# Single cell map of the human ovarian cortex

**DOI:** 10.1101/791343

**Authors:** Magdalena Wagner, Masahito Yoshihara, Iyadh Douagi, Anastasios Damdimopoulos, Sarita Panula, Sophie Petropoulos, Haojiang Lu, Karin Pettersson, Kerstin Palm, Shintaro Katayama, Outi Hovatta, Juha Kere, Fredrik Lanner, Pauliina Damdimopoulou

## Abstract

The human ovary orchestrates sex hormone production and undergoes monthly structural changes to release mature oocytes. The outer lining of the ovary (cortex) has a key role in defining fertility in women as it harbors the ovarian reserve. It has been postulated that putative oogonial stem cells exist in the ovarian cortex and that these can be captured by DDX4 antibody isolation. We analysed on a single cell level the transcriptome and cell surface antigen profiles of over 24,000 cells from high quality ovarian cortex samples from 21 patients. Our single cell mapping reveals transcriptional profiles of six main cell types; oocytes, granulosa cells, immune cells, endothelial cells, perivascular cells, and stromal cells. Cells captured by DDX4 antibody are perivascular cells, not oogonial stem cells. Our data does not support the existence of germline stem cells in adult human ovaries thereby reinforcing the dogma of a limited ovarian reserve.

## INTRODUCTION

The ovaries have two main functions; providing mature and developmentally competent female germ cells oocytes, and producing hormones to support the female phenotype and pregnancy. In contrast to male germ cells spermatozoa that are continuously produced throughout adulthood, oocytes are formed already during fetal life. In the female fetus, primordial germ cells (PGCs, also called fetal germ cells, FGCs) proliferate and migrate to the fetal gonads where they become oogonia [1]. Oogonia proliferate and form cysts that are broken down by intersecting somatic cells into primordial follicles where meiosis is initiated. The oocytes within the primordial follicles remain dormant and arrested in the first meiotic division until the re-activation of follicles in the postnatal ovary [2-4]. The stock of dormant primordial follicles in the postnatal ovary, which reside within the *ca* 1 mm thick ovarian cortex, determines the fertile life span of the woman [5, 6]. Upon activation, they mature in the process called folliculogenesis while migrating towards the inner part of the ovary, medulla [4, 7]. When the reserve of follicles is consumed, menopause commences and the woman cannot naturally conceive anymore. One of the most enigmatic and debated cell types in the ovarian cortex is the oogonial stem cell (OSC). Although it is well understood that women exhaust their ovarian reserve and enter menopause at the age 50, on average [6, 8], there are reports suggesting that adult cortex contains an OSC population stemming from FGCs that is capable of producing new oocytes [9]. These OSCs have been isolated from human ovarian tissue using fluorescence activated cell sorting (FACS) and an antibody that targets the C-terminal domain of DDX4 protein [10]. DDX4 (a.k.a. VASA) is a cytoplasmic RNA helicase specifically expressed in the germline and commonly used as a marker for germ cells including oocytes in adult ovaries [11-13]. In contrast to all other types of germline cells, OSCs have been suggested to express DDX4 on their cell membrane enabling the isolation *via* FACS [9]. Isolated OSCs were shown to express germline markers (e.g. *DAZL, PRDM1*, and *IFITM3*), proliferate *in vitro* and give rise to oocytes in a xeno-transplantation model [9]. The biological significance of these cells is poorly understood. In addition, many laboratories have not been able to repeat these findings and confirm the existence of OSCs [14-16].

Recently, somatic cells from the inner part of human ovaries have been characterized by single cell sequencing [17]. Here, we complete the map of cell types in human ovaries by providing the first extensive characterization of germ cells and somatic cells in the outer layer of human ovaries, *i*.*e*. in the cortex. We used tissue samples provided by 21 patients with proven healthy follicles. Our results demonstrate six main cell types, but cannot provide support to the existence of OSCs. This dataset will be a valuable tool for studying the role of specific cell populations in ovarian biology, dissecting causes of infertility, and developing novel assisted reproductive technologies or even contraceptives.

## RESULTS

### Study setup

We used single cell RNA-sequencing (scRNA-seq) and surface marker screening to determine transcriptomes and cell surface proteomes of cells present in ovarian cortex (Fig. 1). We used tissue from altogether 21 Caesarean section (C-sec, N=5) and gender reassignment surgery patients (GRPs, N=16) to have a comprehensive coverage of individuals and commonly used donor types. All tissue samples were validated to contain healthy follicles (Fig. 1). In order to relate the findings to OSCs, we marked the cells with DDX4 antibodies (Ab) in all experiments; the scRNA-seq experiments were done on uncultured, unsorted cells and sorted DDX4 Ab+ and Ab-cell fractions as well as on cultured, sorted DDX4 Ab+ and Ab-cells. The surface marker screen included DDX4 Ab as an additional marker. Data analysis included validation of markers by immunostaining of ovarian samples, and merging the data with publicly available datasets of cells in the inner part of human ovaries [17] as well as fetal ovaries containing FGCs [18].

**Figure 1.**
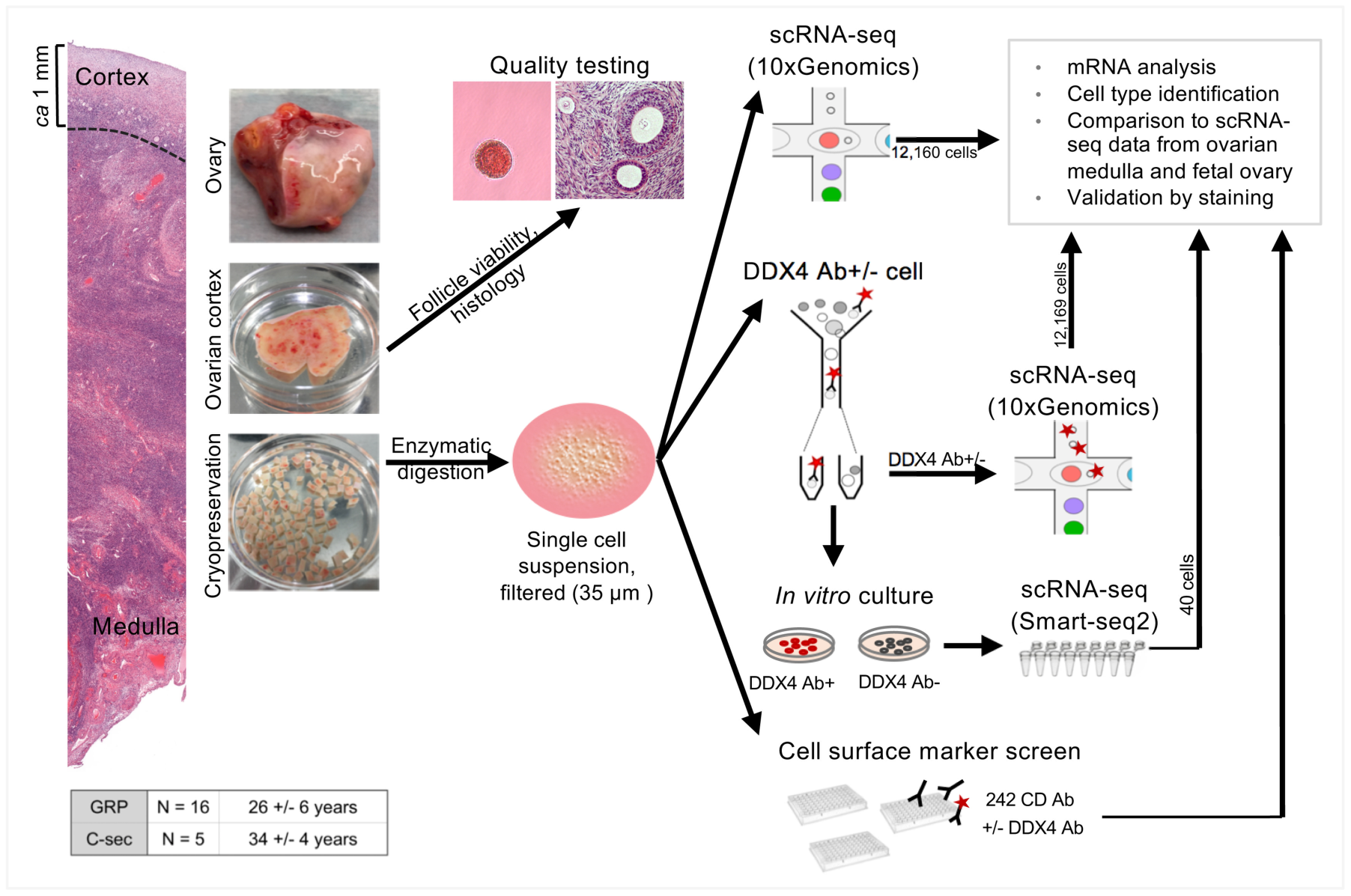
Study design and workflow. Schematic representation of the study. In total, ovarian cortical tissue from 16 GRPs (mean age 25 y) and five C-sec patients (mean age 34 y) was used. Ovaries were trimmed upon tissue receival to separate the cortex. Cortex was cut into pieces, the quality was assessed via follicle viability assay and general histology, and the sample was cryopreserved. Thawed ovarian cortex was enzymatically digested into single cell suspension, filtered, and subjected either uncultured to single cell transcriptome and surface proteome profiling using 10xGenomics technology and cell surface marker screening panel, respectively, or cultured under OSC conditions and subjected to single cell transcriptome analysis using Smart-seq2. In all assays, cells were stained with DDX4 Ab in order to relate the findings to DDX4 Ab+ (suggested OSCs) and Ab-ovarian cell populations. In final data analysis, cortical cell populations were identified, related to cultured and freshly sorted DDX4 Ab+ and Ab-cells, and compared to cells in the inner part of human ovaries [17] and fetal ovarian cell populations [18]. Identified key markers were validated with immunostainings. Ab, Antibody; C-sec, Cesarean section; DDX4, DEAD-Box Helicase 4; GRP, gender reassignment patient; scRNA-seq, single cell RNA-sequencing.

### ScRNA-seq reveals six main cell populations in human ovarian cortex

Ovarian tissue was obtained from one GRP (20 y) and three C-sec patients (28 y, 32 y, 36 y) for unsorted single cell profiling. The cortex containing the primordial follicles was carefully separated from medulla and processed to two independent libraries that were sequenced in parallel (Fig. 1, Supplementary Fig. 1a). After quality control and filtering (see Methods), 12,160 cells were available for cell type characterization (Fig. 1). Uniform Manifold Approximation and Projection (UMAP) analysis revealed six main cell clusters and a similar contribution of cells from GRP and C-sec patients to all clusters (Fig. 2a and Supplementary Fig. 1b). Cell type identities were assigned using gene expression analysis (Fig. 2b, Supplementary Table 1). Uniquely expressed genes revealed distinct cell clusters (Fig. 2c). Signature genes were identified for each cluster and their expression score is shown as violin plots (Fig. 2d). The smallest cluster expressed the typical oocyte markers *GDF9, ZP3, FIGLA* and *OOSP2* (Fig. 2d, Supplementary Table 1). The immune cell cluster expressed tissue resident immune cell markers (e.g. *CD69* and *ITGB2*) as well as mixed gene signatures of T-cells (*CD2, CD3G*, and *CD8A*) and antigen presenting cells, such as *CD14, HLA-DRA, B2M* and *HLA-DQB1*. Endothelial cells could be identified based on the expression of strong endothelial markers such as *VE-cadherin* (*CDH5*) and *VWF*. Granulosa cells were identified based on the expression of typical granulosa cell markers *FOXL2* and *Anti-Müllerian Hormone* (*AMH*). Around 10% of ovarian cortex cells were annotated as perivascular cells expressing *MYH11, MCAM, RGS5, RERGL*, and *TAGLN*, with enriched gene signatures associated with both pericytes and smooth muscle cells [19]. The majority of cells (83%) were broadly classified as stroma. There were no striking differences in gene expression between stromal cells but rather a shared expression of diverse mesodermal lineage markers (e.g. *PDGFRA* and *DCN*) and extracellular matrix proteins (e.g. *COL1A1* and *COL6A1*) (Fig. 2d, Supplementary Table 1).

**Figure 2.**
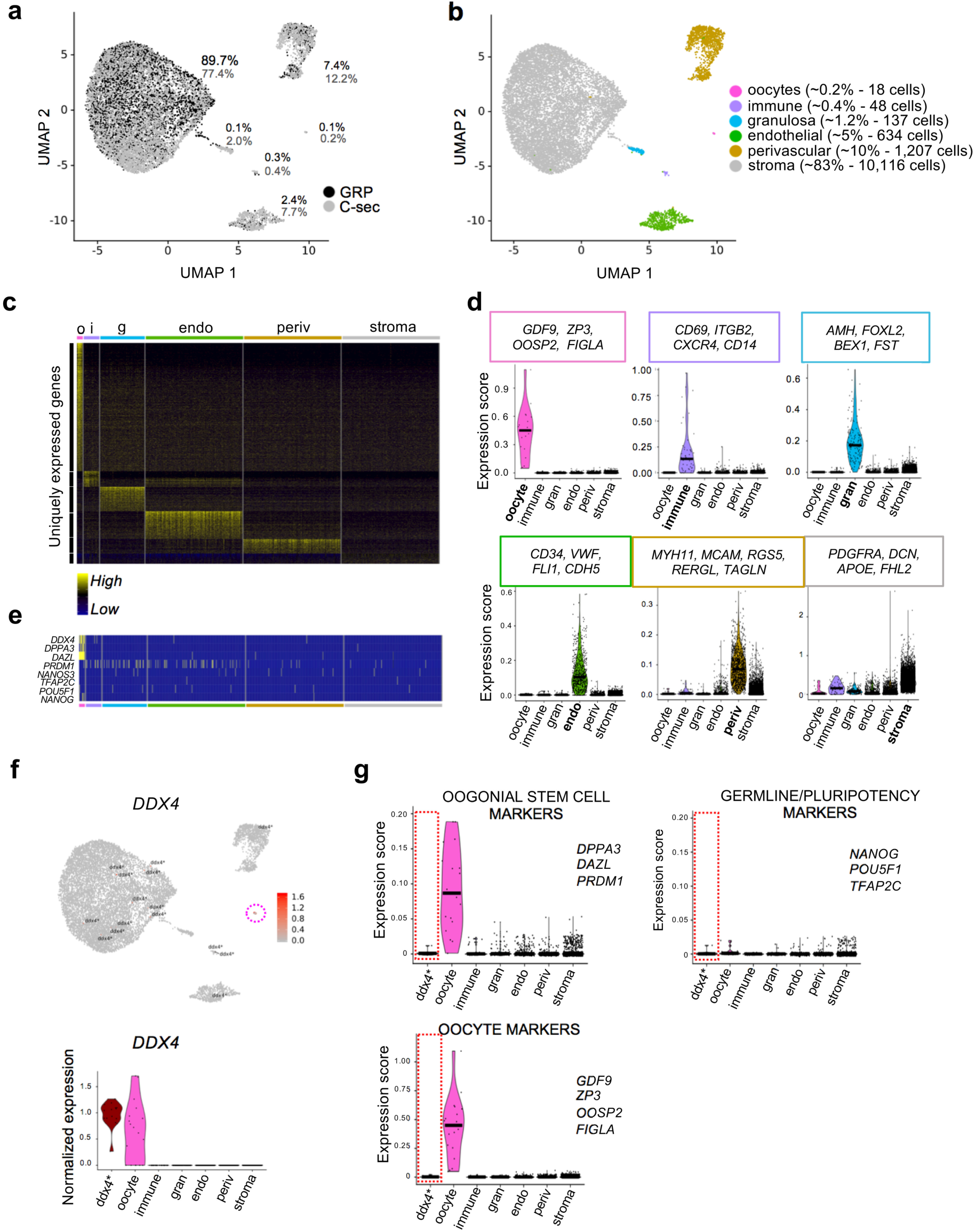
Transcriptome analysis of unsorted ovarian cortex cells. Ovarian cortical cells from GRP and C-sec patients were sequenced on a single cell level. **a**, After quality control and filtration, 5,715 GRP cells (black) and 6,445 C-sec cells (grey) were further analysed. UMAP analysis revealed six main clusters, and both GRP and C-sec cells contributed to all clusters to a similar extent. **b**, Main cell cluster identities were identified with the help of gene expression analysis. **c**, Heatmap showing uniquely expressed genes per cluster. Clusters were downsampled to max. 300 cells per cluster for better visualization. **d**, The color-coded violin plots show the expression scores of signature genes. **e**, Heatmap showing representative marker genes of OSCs and pluripotent cells by cell cluster. **f**, Feature plot displaying all cells expressing *DDX4* transcript. Expression is mainly localized to the oocyte cluster (red circle) and dispersedly in 15 cells from other clusters (labeled with ddx4* for clarity). The 15 somatic cells expressing *DDX4* were manually selected to form their own cluster (ddx4* in violin plot). **g**, Violin plots displaying the expression scores of reported OSC marker genes (*DPPA3, DAZL, PRDM1*), germline and/or pluripotency-associated genes (*POU5F1, NANOG, TFAP2C*) and oocyte marker genes (*GDF9, ZP3, OOSP2, FIGLA)* across all cell clusters. ddx4* cluster does not show differential expression of these markers. The horizontal bars in violin plots indicate median gene expression per cluster. OSC, Oogonial Stem Cell; UMAP, Uniform Manifold Approximation and Projection.

In order to obtain a complete map of cell types in adult human ovaries, we integrated our data with the recently published single cell profiling of antral follicles and tissue fragments manually selected from the inner part of human ovary [17]. While oocytes could not be detected in this dataset [17], most reported somatic cell types overlapped with the cortex cell types identified here including immune cells, perivascular (or smooth muscle) cells and endothelial cells (Supplementary Fig. 1c and d). Interestingly, the cortical granulosa cells clustered separately from the various granulosa cell types found in the antral follicles while the large antral follicle associated theca/stroma cell cluster overlapped with our cortical stroma cluster (Supplementary Fig 1c and d). Collectively, this merged cell map demonstrates that similar vasculature and immune cell clusters can be found throughout ovaries; that there is a close relationship between theca cells and the general stromal cells; and that the granulosa cells are different in the cortical part of the ovary harboring the primordial follicles compared to the inner part containing growing antral follicles.

We next investigated the presence of OSCs in the cortex by studying the expression of reported OSC markers [9] as well as germline stem cell and pluripotency markers. The OSC markers *DDX4, DAZL*, and *DPPA3* were found in the oocyte cluster only (Fig. 2e), while *IFITM3* was detected to a varying extent in all cell types (Supplementary Fig. 1e). We noted that there were sporadic cells throughout the somatic clusters that expressed *DDX4* hence possibly being OSCs (Fig. 2e, 2f). We manually pooled these 15 cells to their own cluster, marked as ddx4*, which showed a similar *DDX4* expression level to oocytes (Fig. 2f) but no differential expression of any other reported OSC (*DPPA3, DAZL, PRDM1)*, germline and pluripotency (*POU5F1, NANOG, TFAP2C*) or oocyte marker (*GDF9, ZP3, OOSP2, FIGLA*) when compared to somatic cells not expressing *DDX4* (Fig. 2g and Supplementary Table 1). Highly expressed genes in the ddx4* cluster (Supplementary Table 1) showed no association with a distinct cell type but rather consisted of various abundantly expressed genes in the cortex. This suggested that the *DDX4*-expressing somatic cells are not germline stem cells.

### Ovarian cortical cells isolated with DDX4 antibody are perivascular cells

Although our data indicated that there are no *DDX4*-expressing germline stem cells in human ovarian cortex, others have isolated such cells using polyclonal Ab (ab13840, Abcam) targeting the C-terminal domain of DDX4 protein [9, 10, 20, 21]. We isolated DDX4 Ab positive (DDX4 Ab+) and negative (DDX4 Ab-) cell populations using the same protocol and tissue source as in the previous reports [9, 10]. We first confirmed that only the recommended DDX4 Ab is able to bind and isolate cells from ovarian cortex. The Abcam Ab bound to a small fraction of ovarian cells (5 – 11.5%) while none of the other tested antibodies against the C-terminal domain enriched any cells above the negative controls (Supplementary Fig. 2a). Using the Abcam Ab, we sorted DDX4 Ab+ and DDX4 Ab-cells from three GRPs (20 y, 22 y, 26 y) (Supplementary Fig. 2b). After quality control and filtration, there were 5,479 Ab+ and 6,690 Ab-cells available for analysis (Supplementary Fig. 2c). As the fraction of DDX4 Ab+ cells in this experiment was 7.7% (Supplementary Fig. 2b), the isolation enriched the population by seven-fold. Seven cell populations emerged from the two libraries (Fig. 3a, Supplementary Fig. 2c). Based on analysis of highly expressed genes (Supplementary Table 2), the same cell types were identified in this combined dataset as in unsorted ovarian cortical cells with the exception of immune cells, which were now separated to T-cells and monocytes (Fig. 3b, Supplementary Fig. 2c and d). The majority (82.5%) of the DDX4 Ab+ cells contributed to a specific cell cluster, which was identified as perivascular cells (Fig. 3a, b and Supplementary Table 2). When considering the seven-fold enrichment of Ab+ cells, the DDX4 Ab-cells dominated all other clusters (Fig. 3a, b and Supplementary Fig. 2c). The top 25 highly expressed genes in Ab+ cells included several known perivascular marker genes such as *MCAM, TAGLN* and *ACTA2*. The top 25 highly expressed genes in the Ab-cells included stromal cell markers, such as *DCN* and *PDGFRA* (Supplementary Table 2). Among the perivascular cell cluster, only one cell expressed *DDX4* (Fig. 3c, left). In the whole dataset, there were two cells in the DDX4 Ab+ and 23 cells in the DDX4 Ab-fractions that expressed *DDX4* mRNA (Fig. 3c, right). Eleven of these were oocytes. The somatic *DDX4* expressing cells were pooled (ddx4*, 1 DDX4 Ab+ cell, 11 DDX4 Ab-cells). Ddx4* cells did not express any germline marker except for *IFITM3*, which was in fact expressed in the majority of the ovarian cortex cells (Supplementary Fig. 2f and g). Ddx4* cells did not express any pluripotency or oocyte markers either (Supplementary Fig. 2g and Supplementary Table 2).

**Figure 3.**
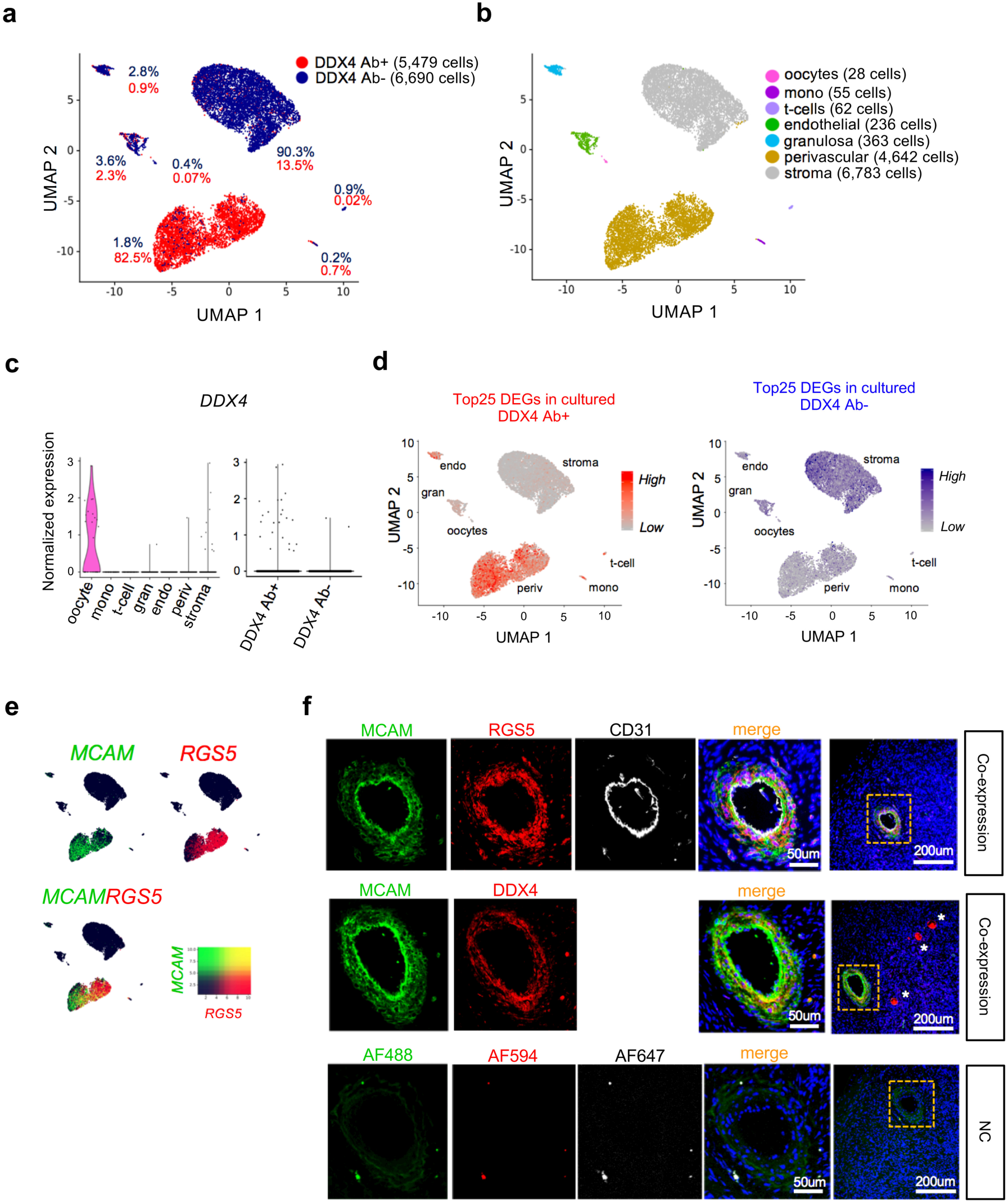
Transcriptome analysis of sorted DDX4 Ab+ and Ab-ovarian cortex cells. Ovarian cortex cells were sorted into DDX4 Ab+ or Ab-populations and sequenced on a single cell level. **a**, After quality control and filtration, 5,479 of Ab+ and 6,690 of Ab-cells were available for downstream analysis. UMAP analysis revealed that 82.5% of the DDX4 Ab+ cells form a distinct cluster while all other clusters mainly consist of DDX4 Ab-cells. **b**, DGE analysis revealed seven main cell types. DDX4 Ab-cells mainly contributed to stroma, endothelial cells, granulosa cells, t-cells, monocytes and oocytes. DDX4 Ab+ cells made up the cluster identified as perivascular cells. **c**, Violin plot showing the expression of *DDX4* among the different clusters (left) as well as among the sorted Ab+ and Ab-populations (right). **d**, Comparison of freshly sorted scRNA-seq data to single cell transcriptome analysis of cultured DDX4 Ab+ and Ab-ovarian cells. Top25 DEGs of cultured DDX4 Ab+ and Ab-cells were superimposed as expression score on to our freshly sequenced DDX4 Ab+ and Ab-cells. The DEGs from cultured DDX4 Ab+ cells (red scale) are highly expressed in the perivascular and endothelial clusters while the DEGs upregulated in cultured DDX4 Ab-cells (blue scale) are not found in these clusters. **e**, Feature plots displaying expression of MCAM (green) and RGS5 (red), both found among top highly expressed genes in DDX4 Ab+ cells, and their co-expression (yellow) on a transcriptional level. Both markers are highly expressed in cells of the perivascular cluster. **f**, Immunostaining of human ovarian tissue sections with MCAM and RGS5, co-stained with the endothelial marker CD31, showed a distinct staining of cells surrounding endothelial cells of blood vessels. In a consecutive section, MCAM is co-expressed in DDX4 Ab+ cells. Oocytes staining brightly positive for DDX4 Ab+ are marked with white asterisks. As negative control, primary antibodies were omitted. DAPI (blue) was used as nuclear counterstain. DEGs, Differentially Expressed Genes; DGE, Differential Gene Expression; MCAM, Melanoma Cell Adhesion Molecule. NC, Negative Control.

OSCs are usually cultured and expanded *in vitro* under specific conditions [9, 10], which could enrich a stem cell population that might go otherwise unnoticed. Therefore, we performed scRNA-seq on FACS-sorted DDX4 Ab+ and Ab-cells cultured under OSC conditions for several weeks. Even after culture, Ab+ and Ab-cells clearly clustered separately (Supplementary Fig. 2h). Highly expressed genes were compared with our uncultured, FACS-sorted ovarian cells. The top 25 highly expressed genes in the cultured DDX4 Ab+ cells associated with the perivascular and endothelial cell clusters whereas the top 25 highly expressed genes in the cultured DDX4 Ab-cells did not (Fig. 3d and Supplementary Table 3). The same results were obtained when the top genes were overlaid with our uncultured, unsorted ovarian cells (Supplementary Fig. 2i). This suggested that *in vitro* culture does not change the identity of the DDX4 Ab+ cells.

We next wanted to verify the localization of DDX4 Ab signal to perivascular cells in human ovarian cortex. We chose two higly expressed markers from the perivascular cluster, *MCAM* and *RGS5* (Fig. 3e), and carried out a co-staining with DDX4. DDX4 was found to be co-expressed with MCAM and RGS5 in the perivascular cells surrounding CD31-positive endothelial cells of blood vessels (Fig. 3f). In the same tissue section, oocytes within pre-antral follicles stained brightly for the DDX4 antibody but not for the perivascular markers (Fig. 3f last column, asterisks).

In order to study the identity of the DDX4 Ab+ cells using an independent approach, we carried out an extensive human cell surface marker profiling of ovarian cells co-stained with DDX4. Fourty-three of the 242 surface markers present in the screen were found to be consistently expressed on ovarian cortex cells (Fig. 4a and Supplementary Table 4). Of these, seven (CD9; CD39; CD44; CD49a; CD49c; CD144; and CD146, also known as MCAM) were brightly expressed on DDX4 Ab+ cells when compared to the DDX4 Ab-cell population, whereas seven (CD26; CD49e; CD54; CD55; CD62e; CD105; CD200) were weakly expressed on DDX4 Ab+ cells (Fig. 4a). Both positive and negative surface marker sets were used to generate expression scores and visualized in feature plots to associate them to cell clusters in our scRNA-seq datasets. Markers expressed on DDX4 Ab+ cells were again associated to the perivascular and endothelial cell clusters, whereas the surface markers absent from DDX4 Ab+ cells were not (Fig. 4b). MCAM and CD9, two markers of DDX4 Ab+ cells identified in the surface marker screen (Fig. 4a) and scRNA-seq analysis (Supplementary Table 2), were validated *via* immunostainings to localize to small blood vessels in the cortex (Supplementary Fig. 3). We next studied the extent to which DDX4 Ab stained perivascular cells. Altogether, 3.67% of the ovarian cells were double positive for the perivascular markers MCAM/CD9, and 80% of them were found to be also positive to DDX4 leading to 2.94% of live cells in ovarian cortex being MCAM+/CD9+/DDX4+ (Fig. 4c). Immunostaining of unfiltered digested ovarian cortex cells likewise showed that all cells of a blood vessel were positive for DDX4 Ab with most of them also co-expressing MCAM and/or CD9 (Fig. 4d). This suggests that the majority of vascular cells are immunopositive for the DDX4 Ab.

**Figure 4.**
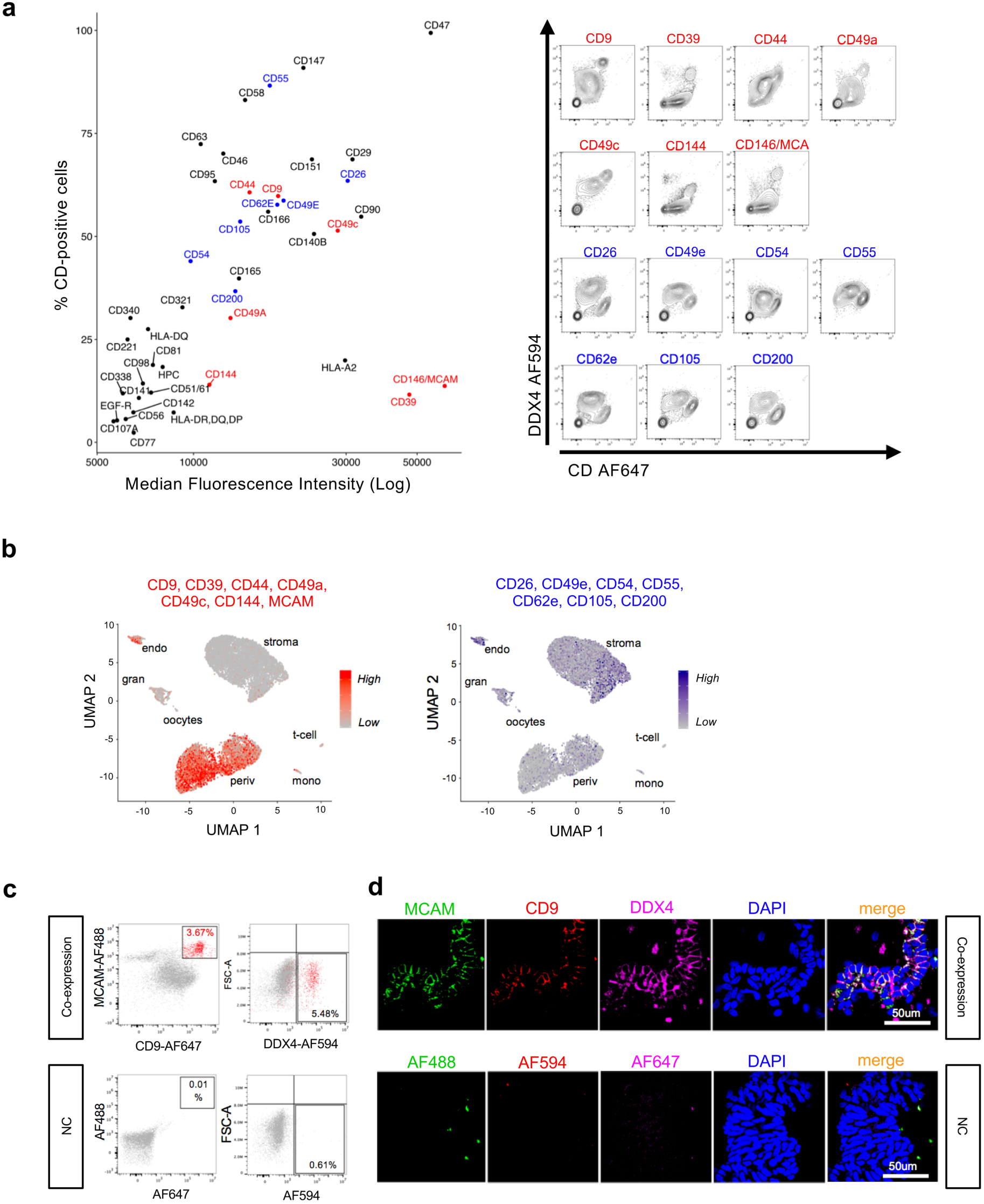
Screening of ovarian cortex cells for surface proteins. The surface of ovarian tissue cells was screened for the expression of 242 CD markers by FC. **a**, Dot plot showing the log median fluorescence intensity (MFI, x-axis) and the percentage of CD-positive cells (y-axis) for those 43 CD markers that were repeatedly found to be present on ovarian tissue cells (left, CD marker expression values of repeat 3 are shown as representatives). When co-stained with DDX4 Ab, seven of the markers were highly expressed (red) and seven were absent or lowly expressed (blue) on DDX4 Ab+ cells when compared to the DDX4 Ab-cell population. Co-expression of DDX4 and the 14 markers identified as high or low in DDX4 Ab+ cells is shown in representative contour plots (right). X-axes display intensity of CD marker signal, y-axes display intensity of DDX4 Ab signal. **b**, Feature plots showing expression scores of the seven cell surface markers highly expressed in DDX4 Ab+ cells (red scale) and the seven markers lowly expressed in DDX4 Ab+ cells (blue scale) overlaid with our sorted scRNA-seq data. While markers that were found to be highly expressed in DDX4 Ab+ cells on a protein level are mainly expressed in perivascular cluster, CD markers lowly expressed in DDX4 Ab+ cells show a more disperse expression pattern. **c**, FACS dot plots showing the co-expression of MCAM and CD9, two markers brightly expressed on DDX4 Ab+ cells, in regard to DDX4 Ab+ and Ab-cell populations. Around 3.67% of ovarian cortex cells were double positive for MCAM and CD9 (in red) and 5.48% are DDX4 Ab+. Around 80% of MCAM/CD9 double positive cells co-stain with DDX4 Ab (2.94% of triple positive MCAM/CD9/DDX4 cells in total). **d**, Unfiltered lysate of ovarian cortex cells cytospinned and stained for MCAM (green), CD9 (red), and DDX4 (magenta) showing co-expression of all three markers on blood vessels. As negative control, primary antibodies were omitted. DAPI (blue) was used as nuclear counterstain. FACS, Fluorescence Activated Cell Sorting; FC, Flow Cytometry; MFI, Median Fluorescence Intensity.

### Absence of germline stem cells in the adult human ovarian cortex

To further investigate the possible existence of germline stem cells in ovarian cortex, regardless of immunopositivity for DDX4 Ab, we compared our data to a recently reported dataset of human fetal ovaries consisting of single cell transcriptomes of fetal germ cells (FGCs) and somatic cells [18]. In these data, FGCs were classified into mitotic FGCs, retinoic acid (RA) responsive FGCs, meiotic FGCs and oogonia, whereas granulosa cells were divided into week 7 – 10, week 10 – 20, and week 20 – 26 granulosa cells [18]. Due to a bias caused by differences in cell numbers between the datasets (1,123 fetal cells in [18] *vs*. 24,329 adult cells in our data), fetal sequencing data were integrated with the individual adult scRNA-seq datasets, resulting in four separate analyses (fetal/C-Sec, fetal/GRP, fetal/DDX4 Ab+, and fetal/DDX4 Ab-) all yielding similar results. A representative dataset (fetal/C-Sec) is shown in Fig. 5a, b. The integrated dataset clustered to nine separate cell types with differing relative contributions by fetal and adult cells (Fig. 5a). Cell cluster identities were attributed based on gene expression analysis (Fig. 5b). FGCs of different stages formed separate clusters that were dominated by fetal cells, whereas fetal oogonia clustered together with adult oocytes. Similarly, fetal somatic cells annotated as endothelial cells clustered together with the adult endothelial cells (Fig. 5a-c). The week 20 – 26 granulosa cells fell into the same cluster with adult granulosa cells, whereas the majority of the earlier stages clustered with adult stroma cells (Fig. 5c and Supplementary Fig. 4a). While none of the 24,329 adult cells sequenced clustered with the meiotic and RA responsive FGCs, four cells clustered with the mitotic FGCs (one cell of C-sec sample, one cell of DDX4 Ab+ sample and two cells of DDX4 Ab-sample) (Fig. 5c and Supplementary Fig. 4a) suggesting that they could potentially be germline stem cells that remain in adult ovaries. The transcription profile of these four cells was investigated further. Fetal mitotic FGCs showed a high expression of the pluripotency genes *PRDM14, POU5F1, NANOG*, and *LIN28A*, but none of these markers were found in the adult cells (Fig. 5d). Consistently, germline marker genes expressed in RA responsive and meiotic FGCs (*TFAP2C, STAR8, SYCP3, NANOS2, NANOS3, DAZL, DPPA3, PRDM1, DDX4, SALL4*, and *BOLL*) as well as oocyte marker genes expressed in fetal oogonia and adult oocytes (*GDF9, ZP3, FIGLA, OOSP2, LIN28, TUBB8*) were not detected in the four adult cells (Fig. 5d). Based on the gene expression profile of the four adult cells and their previous clustering behavior in adult dataset analysis (Fig. 2b and 3b), these cells are considered to be granulosa, perivascular and stroma cells.

**Figure 5.**
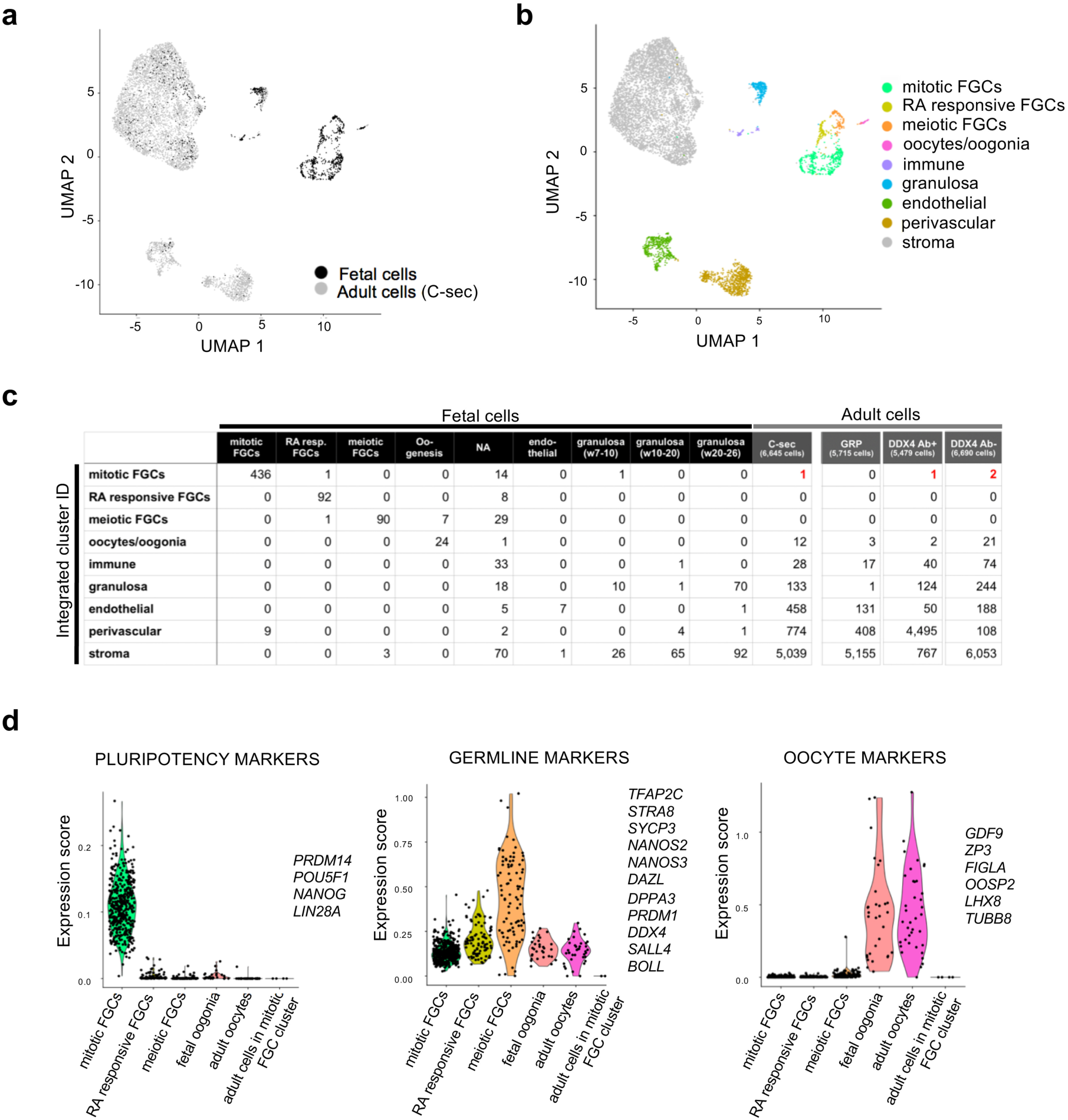
Investigation of potential germline stem cells in the adult human ovarian cortex. Datasets of unsorted and sorted ovarian cortex cells were used individually for integration with scRNA-seq dataset of human fetal tissue cells [18]. **a**, UMAP plot showing sample origin of fetal (black) and adult cells (C-sec data, grey). **b**, UMAP showing nine clusters representing different cell types annotated based on DGE analysis. The same clusters as in adult tissue were found, as well as three new ones: mitotic fetal germ cells, retinoic acid responsive FGCs and meiotic FGCs. **c**, Table listing cell numbers from adult and fetal data sets per annotated cluster. One cell of C-sec sample, one cell of DDX4 Ab+ sample and two cells of DDX4 Ab-sample clustered with mitotic FGC of the fetal sample (in red). These four cells were manually clustered together and studied further with violin plots. **d**, Violin plots displaying expression scores of known pluripotency, germline, and oocyte markers in the different FGC clusters in addition to adult oocyte cluster and the four adult ovarian cells clustering with mitotic FGCs. FGCs, fetal germ cells; NA, non-annotated.

## DISCUSSION

In this study, we provide the first comprehensive map of cell populations in human ovarian cortex, including both oocytes and their somatic cell niche. We identify oocytes, granulosa cells, immune cells, endothelial cells, perivascular cells and stroma cells but do not find a germline stem cell population.

A concern in ovary research is the type of tissue available for research as healthy ovaries from young women are never surgically removed without severe indications. C-sec biopsies are small, GRP ovaries influenced by androgen treatment, and fertility preservation samples affected by the diagnosis and treatments. In addition, patient age has a major impact on follicle density and quality [6, 22]. Typically, GRPs are young and have a high follicle density. Although all GRPs undergo androgen-therapy prior to oophorectomy, their follicles do not differ in distribution, number and quality from untreated women [23], and they can achieve pregnancy after discontinued hormone treatment [24]. In support of this, our data show that ovaries from GRPs have the same cell composition as tissue from C-sec patients. All tissue samples in our study were verified to have good quality and viable follicles, and the samples were handled according to protocols used in clinical fertility preservation [25, 26].

We succeeded in identifying oocytes from the cortical tissue samples. The oocytes likely stem from primordial follicles of the ovarian reserve that resides in the cortex [27]. The digested cell suspension underwent several filtering steps to exclude structures larger than 35 µm of size prior to analysis. This protocol likely excludes oocytes of growing follicle stages as oocytes from primordial follicles are around 35 µm in size compared to primary follicle oocytes that already measure 40 µm [28]. In addition, the oocytes in our data clustered together with fetal oogonia, suggesting that they represent the earliest developmental stage of an oocyte, and stem from the primordial follicles established in the fetal ovaries. Lastly, the expression profile of the detected oocytes lacks markers of oocytes from antral follicles, such as *NTF4, GPD1*, and *LCP2* [13]. The granulosa cells detected in our dataset also appear to stem from very early follicles as they clustered together with fetal week 20 - 26 granulosa cells [18] and lacked known markers of antral granulosa cells such as *CYP19A1* and *FSHR*. In addition, our granulosa cells clustered separate from the ones isolated from antral growing follicles from the inner part of ovaries [17]. Primordial follicle granulosa cells control follicle activation [29] and our data can help isolate these cells for further *in vitro* functional studies.

Ovaries undergo constant remodeling due to folliculogenesis and ovulation, and immune cells have been reported to take part in both processes [30-32]. To further assure proper follicle maturation and restructuring of ovarian tissue, intense formation of new blood vessels composed of endothelial and perivascular cells is taking place in the adult human ovary [33, 34]. Our data are consistent with the known cell types present in ovaries [17], however precise proportions have not been previously reported. In our dataset, macrophages and T-cells make up around 0.5% of the ovarian cortex, endothelial cells around 5% and perivascular cells around 10%.

Steroidogenesis carried out by gonadotropin sensitive theca and granulosa cells of growing follicles is an essential part of ovarian function. Theca cells are acquired during secondary follicle stage and they express *STAR* (*Steroidogenic Acute Regulatory protein*) and *CYP11A1* (*Cholesterol Sidechain Cleavage Enzyme*) enabling them to synthesize androgens [35]. Androgens are further converted to estrogens by CYP19A1 in granulosa cells [35]. We do not detect theca cells in our dataset or other cells with coherent signatures of gonadotropin responsiveness or steroidogenic activity, which is in agreement with known ovarian biology; the cortex contains predominantly primordial follicles that are independent of gonadotropins and steroidogenically inactive. However, some cells in the stroma cluster expressed *STAR* and *CYP11A1*, allowing to hypothesize about a possible presence of early theca cell precursors [36]. The close relationship between theca cells and stromal cells is further reflected by the difficulty to separate theca and stromal cell clusters even in large secondary follicles with visible theca cell layers growth [17]. Identification of factors that trigger theca cell recruitment and markers to identify the early theca will be needed to understand how this cell type relates to general stromal cells in cortex and medulla.

Some medical treatments such as chemotherapy lead to early exhaustion of ovarian reserve and premature ovarian insufficiency [37]. Some patients can benefit from fertility preservation via cryopreservation of ovarian cortical tissue that can be later transplanted for achieving pregnancy [38]. Reports of OSCs have given hope of new assisted reproductive technologies based on cells isolated from ovarian tissue with DDX4 Ab [39-43], although some reports [14-16] and clinical studies [44] contradict the existence of OSCs and their clinical benefit. Our data suggest that there are no OSCs in ovarian cortex, and that the DDX4 Ab+ cells that have been used to treat infertility in women [41] have in fact been perivascular cells. Interestingly, we note that oocytes can be found among the FACS-sorted cells. The presence of oocytes in the DDX4 Ab+ cell fraction could help explain the observations suggesting that DDX4 Ab+ cells can give rise to oocytes [9]. Considering the clinical importance of OSCs, we also addressed their existence using an unbiased approach by merging our data with human FGCs, the suggested origin of OSCs in adult ovaries. None of the studied >24,000 cells expressed an FGC profile, showing that no pre-meiotic fetal germ cells remain in adult ovaries. This observation is in line with the known limited reproductive life span of women, ending in menopause when the cortical follicle pool is exhausted [6].

The FACS-based isolation protocol for OSCs that targets the C-terminal domain of DDX4 protein by a distinct antibody (ab13840, Abcam) [10] has been criticized earlier. DDX4 is an RNA helicase found in the cytoplasm during germ cell development [11] and its expression is maintained at a high level in oocytes [12]. Considering the function of DDX4 in RNA metabolism [45, 46], OSC isolation relying on membrane localization of DDX4 was questioned. Indeed, experimental [47] and *in silico* data [14] argue against an extracellular localization of the DDX4 C-terminus. Despite this, several groups have been able to isolate a varying amount of cells from the ovarian cortex using this antibody, ranging from 1.7 – 42.7% [9, 15, 16, 20, 48]. In our experiments, the population size varies from 5 – 11.5%. The variation could be explained by the organization of the vasculature in the ovary. The initial manual trimming of the poorly vascularized ovarian cortex can lead to contamination with the highly vascularized medulla (see Supplementary Fig. 1a), which could affect the amount of blood vessels and hence perivascular cells present in the sample. In addition, vascular density inversely correlates with follicle density in the cortex [36], and this could lead to varying population size of DDX4 Ab+ cells. Our data also explain how DDX4 Ab+ “OSCs” have even been isolated from extragonadal tissues such as liver and kidney [15]. As all tissues contain blood vessels, perivascular cells are to be expected. We further show that no other Ab targeting DDX4 C-terminus can bind to cells in ovarian cortex, and that the Abcam Ab stains blood vessels (and oocytes) in ovary tissue sections. We conclude that the DDX4 Ab (ab13840, Abcam) recognizes an epitope specifically expressed on perivascular cells.

Perivascular cells include pericytes and smooth muscle cells that originate from a common progenitor and express distinct sets of markers [19]. Depending on the type of blood vessel (fine capillary or big artery) one or both cell types can be found. Interestingly, tissue resident pericytes have been suggested to contribute to the regeneration of various tissues including endometrium due to their stem cell potential [49]. In this study, DDX4 Ab+ cells identified as perivascular cells express markers of pericytes and smooth muscle cells, suggesting that the epitope recognized by DDX4 Ab is expressed on both cell types. The role of perivascular cells in monthly ovarian regeneration should be addressed in future studies.

Altogether, our data suggest that adult human ovarian cortex consists of six main cell types and do not harbor germline stem cells. These data can be used i) to isolate and study the specific populations further *in vitro*; ii) to compare these normal cell profiles to those present in different disease conditions to understand causes of infertility; and iii) to further develop artificial ovaries and other novel assisted reproductive technologies for infertility.

## METHODS

### Ovarian tissue handling

Use of ovarian tissue in research was approved by Stockholm Region Ethical Review Board (Dnr.2010/549-31/2, Dnr. 2015/798-31/2). Independent clinicians informed the patients about the study, and cortical tissue was biopsied from C-sec patients and whole ovaries collected from GRPs after written informed consent. In the described studies, tissue from 16 GRP (20 – 38 yr) and five C-sec patients (28 – 37 yr) was used. All samples were collected at Karolinska University Hospital and transported in Dulbecco’s phosphate-buffered saline supplemented with calcium, magnesium, glucose and pyruvate (DPBS++++, ThermoFisher Scientific, USA) to the research laboratory for processing. In GRP ovaries, cortex was separated from medulla and trimmed to a thickness of 1 mm using scalpels. C-sec biopsies were small surface cuts (max 5×5×1 mm) and did not require trimming.

All tissue samples were cryopreserved and quality controlled using clinical fertility preservation protocols [25, 26, 50]. For histological evaluation, fresh pieces of cortex were immediately fixed in 4% methanol-free formaldehyde (Thermo Fisher Scientific) and in Bouin’s solution (Sigma-Aldrich, USA). Upon dehydration and paraffin embedding, tissue was sectioned (4 µm) and stained with hematoxylin and eosin (HE), and tissue quality was evaluated. For assessment of follicle viability, cortical tissue was chopped using McIIwain Tissue Chopper (Mickle Laboratory, UK) in digestion medium composed of McCoy’s (GIBCO, Thermo Fisher Scientific) medium containing 1 mg/mL human serum albumin (HSA, Vitrolife, Sweden), 1x GlutaMax (Thermo Fisher Scientific), 1x Insulin-Transferrin solution (GIBCO, Thermo Fisher Scientific), 40 µg/mL Liberase™ (Sigma-Aldrich), 0.4 mg/mL Collagenase IV (GIBCO, Thermo Fisher Scientific), and 0.2 mg/mL DNase I (BioRad, USA). The chopped pieces were then incubated in digestion medium containing Neutral Red (50 mg/mL, Sigma-Aldrich) in +37 °C shaking for 30 – 50 min and the digestion was stopped using termination medium containing McCoy’s + 10% fetal bovine serum (FBS, Thermo Fisher Scientific). The presence of viable (red) follicles was verified under a stereomicroscope (Leica S9D) (Fig. 1). For vitrification of ovarian tissue, our established vitrification protocol was used [51]. Briefly, tissue was cut into pieces of 1×1×1 mm and transferred into vitrification solution containing 40% Ethylene Glycol (Sigma-Aldrich) and 10 mg/mL HSA for 2 min and 3 min. Then, pieces were transferred into cryo-tubes and stored in liquid nitrogen. For thawing, cryo-tubes were slightly opened and equilibrated in room temperature for 30 s. Closed tubes were then placed into a 37 °C water bath for 1.5 min, and the pieces were transferred through three different thawing solutions of decreasing sucrose and increasing HSA concentration for 2 min, 3 min and 5 min. For slow-freezing of ovarian tissue, the standard protocol was followed [26]. In brief, cortex was cut into pieces of a maximum size of 10×10×1 mm and pre-equilibrated in slow-freezing medium containing 7.5% of Ethylene Glycol and 10 mg/mL HSA in DPBS for 30 min shaking on ice. Pieces were transferred into cryo-tubes containing 1 mL of fresh slow-freezing solution and placed into controlled rate freezer (Kryo 360-1.7, Planer PLC, UK). For thawing of slow-frozen tissue, cryo-vials were placed in a 37 °C water bath for 1 – 2 min, transferred into thawing solutions with 10 mg/mL HSA and decreasing sucrose concentrations for 10 min, 10 min, and 10 min.

### Dissociation of human ovarian cortex into single cell suspension

Thawed human ovarian cortical tissue was chopped using scalpels into pieces of approximately 0.3 mm^3^ and enzymatically digested in DMEM/F12 (Thermo Fisher Scientific) containing 5% FBS, 1 mg/mL collagenase IA (Sigma-Aldrich), 50 μg/mL Liberase™ and 1000 U DNase I (Roche, Sigma-Aldrich) in a shaking 37 °C water bath for max. 50 min. Digestion was stopped with medium containing 10% FBS and cell suspension was centrifuged for 7 min on 300 g. Cells were resuspended in DPBS, 2% FBS and passed through a 40 μm cell strainer (VWR, USA), counted and used for subsequent experiments.

### Flow Cytometry (FC) and Fluorescence-Activated Cell Sorting (FACS)

Dissociated ovarian cell suspension was prepared for FC (surface marker screen using seven GRPs and validation of markers using three GRPs) and FACS (scRNA-seq using three GRPs) following the previously published oogonial stem cell isolation protocol [10]. All blocking and antibody incubation steps were carried out for 20 min on 4 °C. After blocking in PBS containing 0.02% bovine serum albumin (BSA, Sigma-Aldrich and Normal Donkey Serum (1:50, EMD Millipore, Germany), samples were washed and stained using primary antibodies (DDX4, ab13840, Abcam; DDX4, SAB1300533, Sigma-Aldrich; DDX4, AP1403b, Abgent; MCAM, AF932, R&D Systems; CD9, ab2215, Abcam) diluted in blocking solution. DDX4 antibody was titrated using concentrations ranging from 5 – 20 μg/mL in 100 μL of cell suspension (max 1×10^6^ cells) and recommended isotype control antibody (polyclonal rabbit IgG, ab171870, Abcam) was used as negative control. A distinct population of DDX4 Ab+ cells were found using a concentration of 10 μg/mL of DDX4 Ab (ab13840) in 100 μL, hence, all subsequent experiments were performed using this concentration (Supplementary Fig 2a). After 20 min, cells were washed and incubated with secondary antibodies diluted in blocking solution: AF488 donkey anti-goat IgG (A11055), AF594 donkey anti-rabbit IgG (A21207), AF647 donkey anti-mouse IgG (A31571, all from Thermo Fisher Scientific). DAPI (0.2 μg/mL, Thermo Fisher Scientific) was added to the final cell suspension as a live/dead marker. Before sorting, cells were transferred into round bottom tubes through a 35 µm nylon mesh (Falcon, Sigma-Aldrich). Negative and positive gates for analyzing and sorting cell populations were set using FMO (fluorescence minus one) controls, isotype controls as well as samples where primary antibody was omitted. The cells were either analysed by FC using CytoFLEX S (Beckman Coulter, USA) equipped with 405, 638, 488 and 561 nm lasers or sorted using a BD FACSAria Fusion (BD Bioscience, USA) equipped with 405, 640, 488, 355 and 561 nm lasers. BD FACSDiva™ software or FlowJo v10 software (Tree Star) was used for downstream analysis and results are presented as representative FACS plots.

### Single cell library preparation and mRNA sequencing (scRNA-seq) of uncultured ovarian cells using 10xGenomics

Two runs of scRNA-seq of dissociated, uncultured ovarian tissue cells were performed. Each run consisted of two samples that were prepared and sequenced in parallel. In the first run, unsorted cortical cells from one GRP and three pooled C-sec patients were analysed. The samples were digested into single cells in parallel and dead cells were removed (dead cell removal kit, MACS, Milteny Biotec, Bergisch-Gladbach, Germany). In the second run, cortex from three GRPs were pooled, digested into single cells, stained with DDX4 antibodies, and sorted into live (DAPI negative) DDX4 Ab+ and DDX4 Ab-cell populations. The cell suspensions were transported on 4°C to the Eukaryotic Single Cell Genomics Facility (ESCG, SciLifeLab, Stockholm, Sweden) and prepared immediately for loading into the 10XGenomics Chromium controller for gem formation. For scRNA-seq library preparation, 10XGenomics v2 (unsorted cells) or v3 (sorted populations) was used according to manufacturer’s instructions and sequenced on a NovaSeq 6000 using a S1 flow cell.

### ScRNA-seq data analysis of uncultured ovarian cells

ScRNA-seq output files were converted using Cell Ranger 2.1.1 (unsorted cells) or 3.0.1 pipeline (sorted cells) and aligned to the hg19 transcriptome using STAR aligner [52]. Analysis of filtered cells was performed in R version 3.5.1 [53] using Seurat suite version 3.0.0 [54, 55]. Details of the web summary of Cell Ranger statistics of unsorted and sorted datasets can be found in Supplementary Table 1 and 2, respectively. For downstream bioinformatics analysis of remaining cells after the initial Cell Ranger filtration based on correctly detected cellular barcodes, genes expressed in at least three cells were kept. In order to exclude potential doublets, cells expressing 200 – 7000 genes, and no more than 25% of mitochondrial genes were kept, resulting in 24,329 cells in total (12,160 unsorted and 12,169 sorted cells). For integration of different datasets (unsorted GRP/C-sec), Seurat’s CCA integration tool was used. The sorted datasets were merged and regressed for batch effects.

All datasets were column-normalized and log-transformed prior to selection of highly variable genes which were used for Principal Component Analysis (PCA). Elbow plot function in Seurat was used to identify most significant PCs that were included when UMAP analysis was performed (dimensionality of 13 in unsorted and 12 in sorted dataset). A resolution of 0.1 and a perplexity of 30 was chosen for analysis. In scRNA-seq analysis of unsorted and sorted datasets, oocytes were selected manually based on marker gene expression analysis using CellSelector function in Seurat. Marker genes discriminating the different clusters were selected among highly expressed genes (p-value < 0.01 and log(fold-change) > 0.25) using Wilcoxon Rank Sum test. Additionally, signature gene expression was calculated as percentage of sum counts of selected genes vs all expressed genes. The percentage ratio is added as feature value to each cell and multiplied by 100.

### *In vitro* culture of DDX4 Ab+ and DDX4 Ab-ovarian tissue cells

Ovarian tissue of two C-sec patients (36 y, 37 y) was pooled before dissociated into single cells and sorted based on DDX4 Ab signal as previously reported [15]. Sorted DDX4 Ab+ and Ab-cell populations were cultured as described earlier [15]. Briefly, after sorting, cell populations were plated onto irradiated mouse embryonic fibroblasts (Applied StemCell, USA) in medium consisting of MEMα, 10% FBS, 1 mM sodium pyruvate, 1 x Glutamax, 1% non-essential amino acids, 0.1 nM 2-mercaptoethanol, 1 x N2-supplement (R&D Systems), 50 U/mL penicillin, 50 ug/mL streptomycin, 10 ng/mL recombinant human leukemia inhibitory factor (Merck Millipore), 1 ng/mL basic fibroblast growth factor (R&D Systems), 40 ng/mL glial cell-derived neurotropic factor (R&D Systems), and 10 ng/mL recombinant human epidermal growth factor (all from Life Technologies if not mentioned otherwise). After passage 3, cultured cells were plated onto MEF-free 0.1% gelatin-coated plates (Sigma-Aldrich). DDX4 Ab+ cells and DDX4 Ab-cells were maintained for seven and six passages, respectively.

### Single cell library preparation and mRNA sequencing (scRNA-seq) of cultured ovarian tissue using Smart-seq2

After culture, DDX4 Ab+ and Ab-cells were randomly picked by mouth pipetting into lysis buffer and cDNA libraries were prepared as described earlier [15]. In brief, whole cell lysates were reversely transcribed using SuperScript II reverse transcriptase (Invitrogen, USA) and resulting PCR products were preamplified using KAPA HiFi HotStart ReadyMix (KAPA Biosystems, USA). Spiked-in samples were sequenced using the TrueSeq dual-index sequencing primers (illumina, USA) according to manufacturer’s instructions on a HiSeq 2000.

### Analysis of scRNA-seq data of *in vitro* cultured ovarian tissue cells

STAR aligner [52] was used to index the hg19 reference genome and align the resulting fastq files. Mapped reads were then counted in annotated genes using featureCounts. The annotations and reference genome were obtained from UCSC Genome Browser [56]. The count table from featureCounts was imported into R/Bioconductor and differential gene expression was performed using the EdgeR package and its general linear models pipeline. For gene expression analysis, genes with at least one count per million in three or more samples were used and normalized using TMM normalization. UMAP dimensionality reduction was performed in R with the uwot package using all genes with at least 1 count per million in three or more samples.

### Integration of fetal scRNA-seq data and adult ovarian medulla scRNA-seq data with ovarian cortex scRNA-seq data

The UMI count data of human fetal scRNA-seq were downloaded from http://github.com/zorrodong/germcell [18]. Female samples expressing 2000 – 9000 genes with no more than 1,100,000 transcripts were kept, resulting in 1,123 cells in total. FindIntegrationAnchors and IntegrateData functions in Seurat [55] were used with dimensionality of 15 – 40 for the integration of these fetal scRNA-seq data with the individual adult scRNA-seq datasets (C-sec, GRP, DDX4 Ab+, and DDX4 Ab-, respectively). The cell cycle scores based on G2/M and S phase markers were calculated by CellCycleScoring function, and these scores were regressed out using ScaleData function to remove the cell cycle effect. Identification of the clusters was determined using the FindClusters function with the resolution which classifies the FGCs into 4 clusters as reported in [18] (resolution of 0.9 – 1.2).

Similary, scRNA-seq data of the inner part of human ovaries, derived from 31 tissue samples of 5 ovaries, were obtained from Gene Expression Omnibus (GEO) using accession number GSE118127 [17]. These data were processed with Seurat (v2.3.0) and scran (v1.12.0) to correct the patient bias by mutual nearest neighbors (MNNs) [57], following the method described in [17]. Finally, 20,676 cells were clustered into 19 clusters as reported, which were then integrated with our 12,610 unsorted ovarian cortex cells, using FindIntegrationAnchors and IntegrateData functions in Seurat (v3.0.0) with dimensionality of 21. Cell cycle effect was removed as described above.

### CD marker screen of human ovarian cortex cells

After dissociation of thawed ovarian cortex tissue into a single cell suspension, cells were stained using the BD Lyoplate™ Screening Panels (BD Biosciences) of 242 monoclonal antibodies following manufacturer’s instructions. Altogether three runs were carried out. In the first run, pooled tissue cells from five GRP ovaries were used for the full plate of 242 CD markers. The subsequent two runs were performed with pooled tissue cells from four and six GRP ovaries, respectively, to verify the expression of the markers identified in the first run. DDX4 co-staining (ab13840, Abcam) was carried out in all runs to identify markers on the surface of DDX4 Ab+ cells. All samples were analysed in a 96-well format on a CytoFlex S flow cytometer (Beckman Coulter). Exclusion of dead cells was ensured by gating on the DAPI-negative cell population using FlowJo v10 software (Tree Star). Results are presented as representative contour plots, percentage of positive live cells and median fluorescence intensity.

### Immunocytochemistry of dissociated ovarian cells

Ovarian tissue from three GRPs was dissociated separately into a bulk cell suspension passed through a 100 μm cell strainer (VWR, USA) to leave vessel structures intact and diluted to a concentration of 50,000 cells in 100 μL of DPBS (ThermoFisher Scientific) containing 2% FBS. Samples were cytospinned onto glass slides (VWR) using Cytocentrifuge (CytoSpin 4, Thermo Fisher Scientific) at 1,200 rpm for 4 min. Slides were dried in room temperature for 2 – 4 hours and fixed with 4% methanol free formaldehyde for 10 min. Cells were permeabilized with 0.3% Triton X-100 (Sigma-Aldrich) in DPBS for 10 min, followed by blocking with in DPBS/ 4% FBS/ 0.1% Tween-20 (P9416, Sigma-Aldrich) for 1 hour. Primary antibodies (DDX4, ab13840, Abcam; CD9, 312102, BioLegend; MCAM, AF932, R&D systems) were diluted in blocking solution and cells were stained overnight at 4 °C, followed by incubation with secondary antibodies for 2 h at room temperature (AF488 donkey anti-mouse IgG, AF555 donkey anti-goat IgG, AF647 donkey anti-rabbit IgG; all from Thermo Fisher Scientific, A21207, A21432, A31573, respectively.). Cell nuclei were counterstained using Hoechst (1:1000, Invitrogen, Thermo Fisher Scientific). Cells were mounted using fluorescent mounting medium (Dako Agilent, USA, CAlifornia) and imaged with Olympus IX81 fluorescence microscope (Carl Zeiss Meditec, Germany) or Nikon spinning disk confocal microscope (Nikon, Germany). Acquired images were analysed using Fiji/ImageJ software v2.0.

### Immunohistochemistry of ovarian tissue sections

Immunohistochemistry was performed on freshly formaldehyde-fixed paraffin-embedded cortex sections (4 μm) of at least two GRPs in technical triplicates. After de-paraffinization and rehydration, antigen retrieval was performed in either 10 mM Sodium Citrate (pH 6) or 10 mM Tris/ 1mM EDTA (pH 9) at 96 °C for 30 min. After blocking in 5% BSA and Normal Donkey Serum, tissue sections were incubated in primary antibody (DDX4, ab13840, Abcam; CD9, 312102, BioLegend; MCAM, AF932, R&D systems, RGS5, ab186799, Abcam; CD31, M0823, Dako) overnight at 4 °C, followed by incubation with secondary antibody for 1 h at room temperature: AF488 donkey anti-mouse IgG, AF555 donkey anti-goat IgG, AF647 donkey anti-rabbit IgG (all from Thermo Fisher Scientific, A21207, A21432, A31573, respectively). Stained tissue sections were mounted in fluorescent mounting media (Dako Agilent) and imaged with Olympus IX81 fluorescence microscope (Carl Zeiss Meditec, Germany) or Nikon ultra fast widefiled microscope (Nikon, Germany). Acquired images were analysed using Fiji/ImageJ software v2.0.

### Data availability

10xGenomics and Smart-seq2 scRNA-seq data have been deposited on ArrayExpress and are available under the accession codes E-MTAB-8381 and MTAB-8403, respectively. ScRNA-seq data analysis was performed using scripts in R and are available upon request.

## Supporting information

Cell Ranger metrics and gene expression analysis of unsorted ovarian cortex cells.

Cell Ranger metrics and gene expression analysis of sorted ovarian cortex cells.

Surface marker screening of human ovarian cortex cells.

Differentially expressed genes in cultured ovarian DDX4 Ab+ and DDX4 Ab- cortex cells.

## ACKNOWLEDGEMENTS

The authors thank all clinicians helping us with patient recruitment and tissue collection as well as all patients donating ovarian tissue to our studies. In particular, the irreplaceable help from Catarina Arnelo and Boel Niklasson is warmly acknowledged. Claus Yding Andersen is thanked for kindly teaching us controlled rate freezing and follicle viability assay. Richelle Björvang and Astrud Tuck are thanked for the help with ovarian tissue handling. Flow cytometry analysis and cell sorting were performed at the MedH Flow Cytometry core facility, Karolinska Institutet, Sweden. Sequencing was performed at the ESCG at Science for Life Laboratory, Stockholm, Sweden. We would like to thank Tarja Schröder and Morphological Phenotype Analysis facility, Karolinska Institutet, Sweden, for H&E stains and tissue scans. Imaging of stained samples was performed at the Live Cell Imaging facility/Nikon Center of Excellence, Karolinska Institutet, Sweden.

This work was supported by the Swedish Research Council (P.D, M.W., F.L.), the Swedish Research Council for Sustainable Development (P.D.), the Swedish Childhood Cancer Foundation (P.D.), Karolinska Institutet PhD student funding (M.W.), the Knut and Alice Wallenberg Foundation (J.K. and F.L.); Wallenberg Academy Fellow, Ragnar Söderberg Foundation, The Ming Wai Lau Centre for Reparative Medicine, Center for Innovative Medicine/SLL (F.L.); the Scandinavia-Japan Sasakawa Foundation, the Japan Eye Bank Association, the Astellas Foundation for Research on Metabolic Disorders, and the Japan Society for the Promotion of Science Overseas Research Fellowships (M.Y.).

## AUTHOR CONTRIBUTIONS

These authors contributed equally: Fredrik Lanner, Pauliina Damdimopoulou.

### Contributions

M.W. carried out the experiments, analysed data, performed sequencing analysis and wrote the paper. M.Y., S.K. and A.D. carried out bioinformatic analyses. I.D. conceived the experiments and sorted cell populations. S.Panula and S.Petropoulos performed scRNA-seq experiment of cultured ovarian tissue cells. H.J. contributed to the surface marker screen verification. K.Palm and K.Pettersson coordinated patient recruitment and carried out surgeries. O.H. conceived the experiments and supervised the project. J.K. contributed to essential resources and helped draft the manuscript. F.L. and P.D. conceived the experiments, analysed data, supervised the project and wrote the paper. All authors contributed to the preparation of the paper and approved the final version.

### Corresponding authors

Corresponding authors: Fredrik Lanner, Pauliina Damdimopoulou.

