## Supplementary Figures for "Single cell map of the human ovarian cortex"

### SUPPLEMENTARY FIGUES

#### Supplementary Figure 1. Cell map of the human ovary.

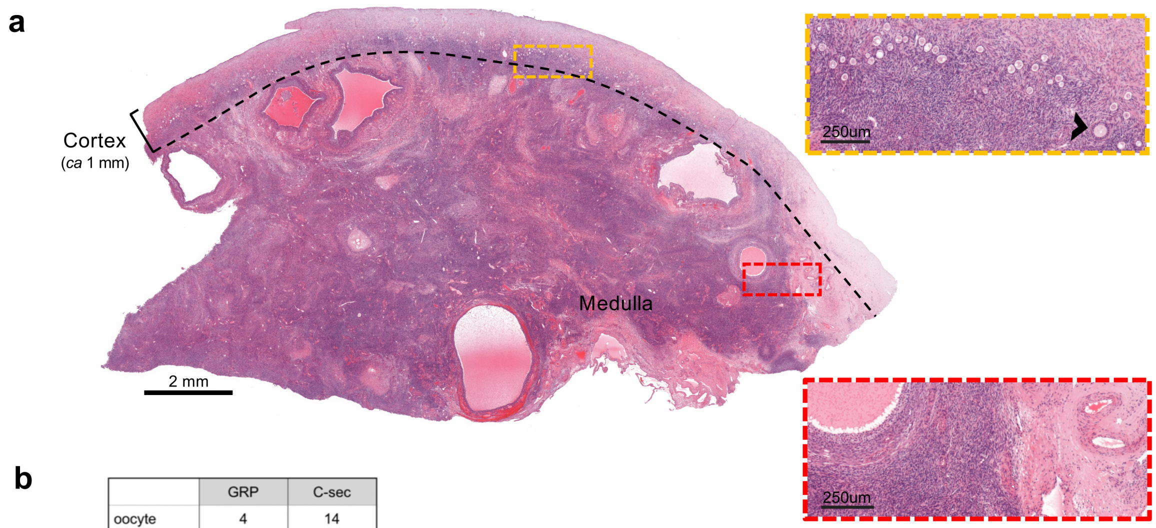


b

| | GRP | C-sec |
|---|---|---|
| oocyte | 4 | 14 |
| immune | 19 | 29 |
| granulosa | 5 | 132 |
| endothelial | 140 | 494 |
| perivascular | 422 | 785 |
| stroma | 5,125 | 4,991 |
| total | 5,715 | 6,445 |

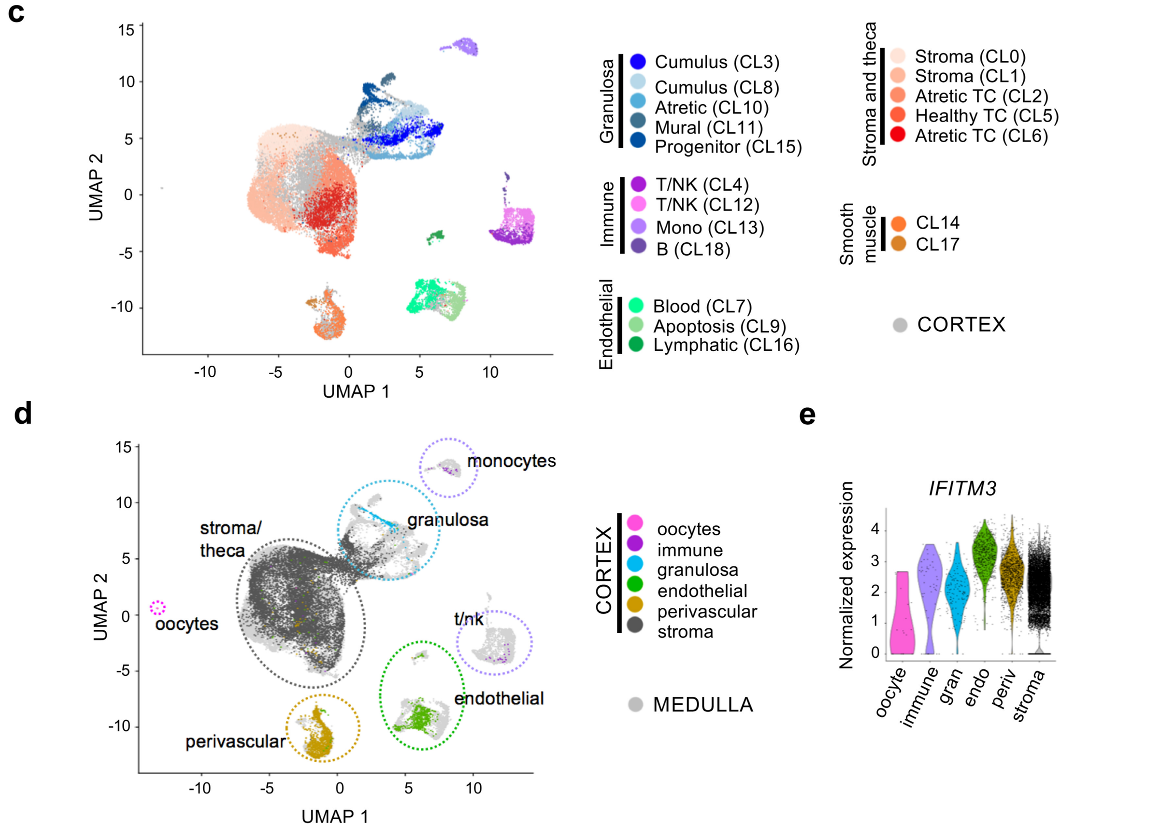


**a**, Representative ovary cross-section of a 26-year old GRP showing the thin cortex layer (*ca* 1 mm, used for fertility preservation) harboring clusters of small primordial follicles that form the ovarian reserve. Beneath the cortex is the inner part of the ovary, medulla, with looser

connective tissue, denser vasculature and larger growing follicles. Yellow rectangle is enlarged to highlight the ovarian reserve composed of primordial follicles resting in the cortex. One early growing (primary) follicle is indicated (arrowhead). Red rectangle is enlarged to highlight the more prominent vasculature in the medulla. **b**, Number of cells contributing to the different cell clusters from GRP and C-sec tissue (in Fig. 2b). **c**, UMAP of merged scRNA-seq data of our unsorted ovarian cortex cells (grey) and the publicly available scRNA-seq dataset of antral follicles and pieces of stroma picked from ovarian medulla (colored cells) [17]. Previous cluster annotations of both datasets are maintained [17]. **d**, Same UMAP as in **c**, showing the contribution of our cortex cells (colored) to previously reported ovarian medulla clusters (grey). All identified cortical cell clusters joined the medullary cell clusters except from the oocytes that were unique to the cortex dataset. **e**, Violin plot showing the expression levels of *IFITM3* in unsorted ovarian cortex cells.
The horizontal bars in violin plots indicate median gene expression per cluster.
CL, Cluster; TC, Theca Cell.

**Supplementary Figure 2. Sorted ovarian DDX4 Ab+ and Ab- cortex cells.**

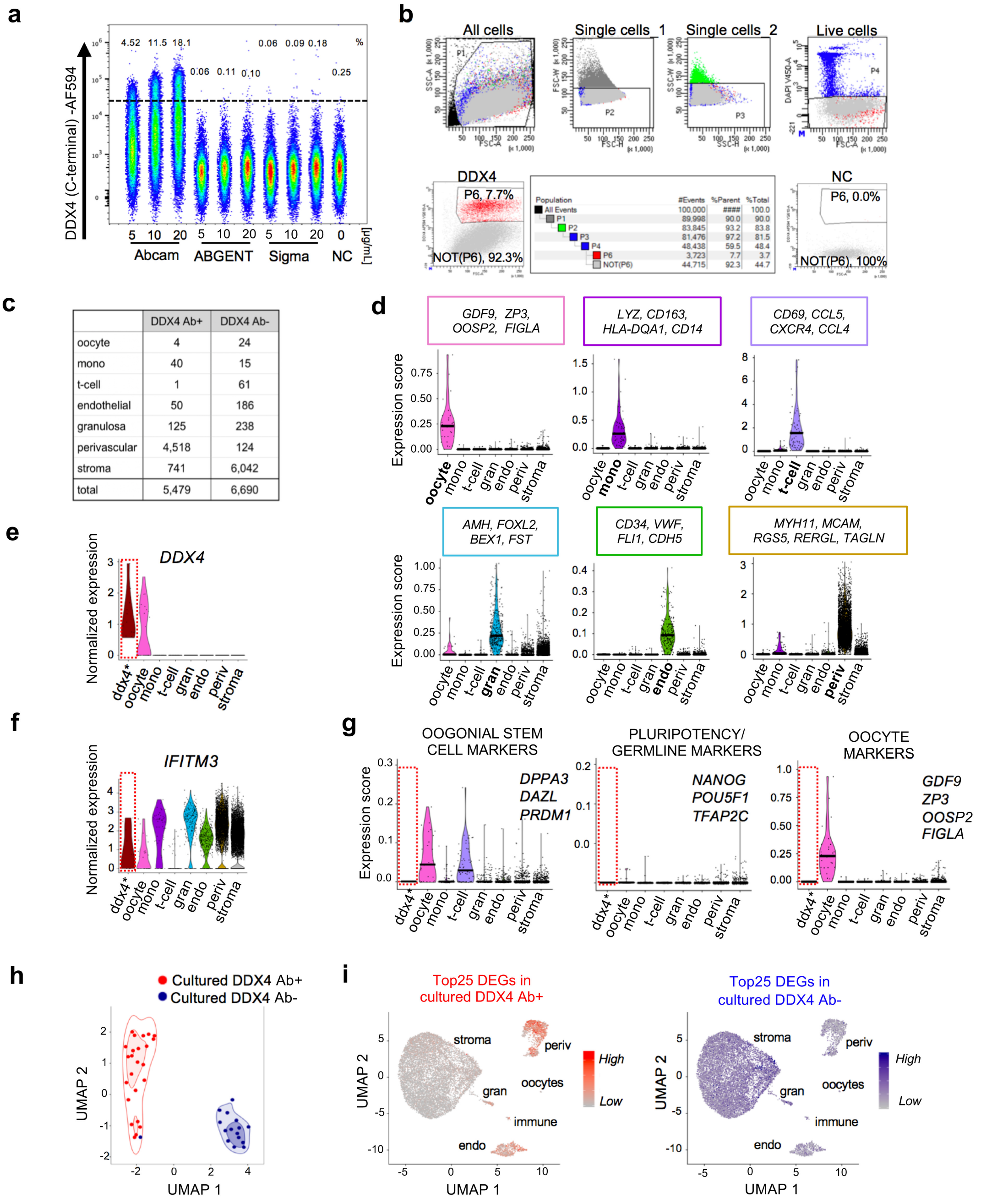


| | DDX4 Ab+ | DDX4 Ab- |
|---|---|---|
| oocyte | 4 | 24 |
| mono | 40 | 15 |
| t-cell | 1 | 61 |
| endothelial | 50 | 186 |
| granulosa | 125 | 238 |
| perivascular | 4,518 | 124 |
| stroma | 741 | 6,042 |
| total | 5,479 | 6,690 |

**a**, FACS plots of three different DDX4 C-terminus antibodies (Abcam, ABGENT, and Sigma) in three different staining concentrations (5, 10 and 20 µg/mL per sample containing max $1x10^{6}$ cells in 100 µL) are shown. The Abcam Ab labeled an increasing percentage of a distinct cell population (4.51% - 18.1%) with increasing concentration while ABGENT and Sigma antibodies did not. Gating of DDX4 Ab+ cells was performed on cells that stained for rabbit polyclonal isotype control Ab. **b**, Representative FACS plots showing the strict gating strategy of dissociated ovarian tissue cells for sorting of DDX4 Ab+ cells for scRNA-seq analysis. After exclusion of debris, single cells were gated based on FSC and SSC properties. Live cells were selected based on DAPI signal. Gating of DDX4 Ab+ cells was based on FSC and distinct Ab signal. Several negative control samples were included in each sort (DAPI only, primary Ab omitted, FMO as well as rabbit polyclonal isotype control, labelled here as NC) to avoid false positive DDX4 Ab+ cells. On average, a DDX4 Ab+ cell population of 5.5 – 11.5% was observed. **c**, Contribution of cells from sorted DDX4 Ab+ and DDX4 Ab- populations to each cluster (in Fig. 3b). **d**, Color-coded violin plots showing expression scores of selected signature genes among the top highly expressed genes defining the cell type of each cluster. Horizontal bars indicate median gene expression per cluster. **e**, Somatic cells expressing *DDX4* (n=12; n=1 in Ab+ population and n=11 in Ab- population) were manually clustered to form a separate cluster (ddx4*). **f**, Violin plot showing the expression levels of *IFITM3* in all sorted ovarian cortex cells. **g**, Violin plots showing OSC (*DPPA3, DAZL, PRDM1*), pluripotency and/or germline (*NANOG, POU5F1, TFAP2C*) and oocyte markers (*GDF9, ZP3, OOSP2, FIGLA*) as expression score. None of the markers were expressed in somatic DDX4 expressing cells (except from *IFITM3)*. **h**, UMAP based on single cell transcriptomes of cultured DDX4 Ab+ and Ab- cells showing that the populations cluster separately even after extended time in cell culture in OSC culture conditions. Contour lines indicate the 2D kernel density for each cell population divided into three bins. **i**, Feature plots showing the expression score of the top25 DEGs of cultured DDX4 Ab+ (red) and DDX4 Ab- cells (blue) in our freshly sequenced unsorted ovarian cells.

Ab, Antibody; DEGs, Differentially Expressed Genes; FACS, Fluorescence Activated Cell Sorting; FMO, Fluorescence Minus One; FSC, Forward Scatter; NC, Negative Control; OSC, Oogonial Stem Cell; PCA, Principal Component Analysis; SSC, Side Scatter.

**Supplementary Figure 3. Validation of markers identified in CD surface marker screen.**

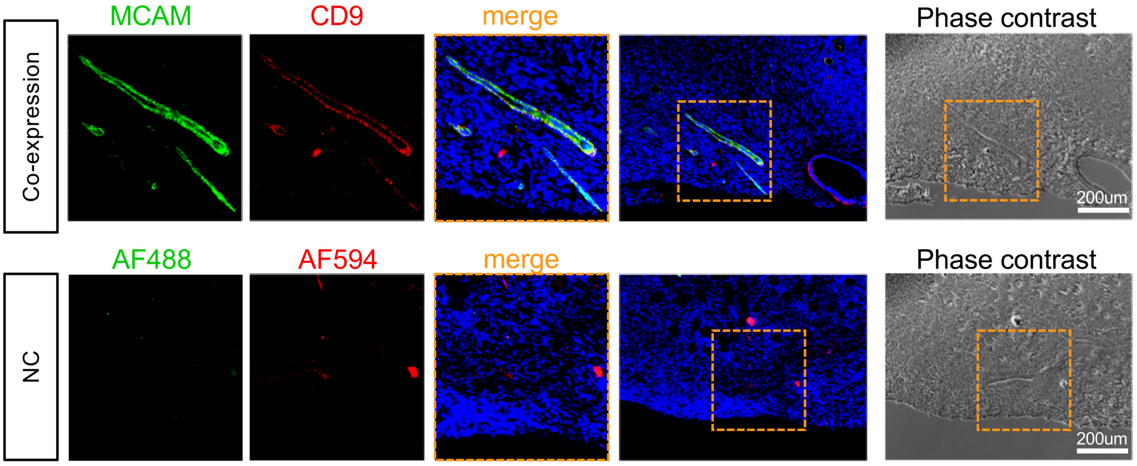


Immunostaining of human ovarian cortical tissue sections from GRP donor showing the expression of MCAM (green) and CD9 (red) on a protein level. DAPI (blue) was used as nuclear counterstain. Co-expression was detected in cells of blood vessels (enlarged). Orange rectangles demarcate the zoomed-in area. As negative control (NC), primary antibodies were omitted.

**Supplementary Figure 4. Germline-like cells in the adult human ovary.**

Integration with unsorted GRP data

Integrated cluster ID

| | mitotic FGCs | RA resp. FGCs | meiotic FGCs | Oo-genesis | NA | endo-thelial | granulosa (w7-10) | granulosa (w10-20) | granulosa (w20-26) | GRP (5,715 cells) |
|---|---|---|---|---|---|---|---|---|---|---|
| mitotic FGCs | 426 | 14 | 0 | 0 | 17 | 0 | 0 | 0 | 0 | 0 |
| RA responsive FGCs | 0 | 78 | 0 | 0 | 8 | 0 | 0 | 0 | 0 | 0 |
| meiotic FGCs | 2 | 2 | 90 | 6 | 27 | 0 | 0 | 0 | 0 | 0 |
| oocytes/oogonia | 0 | 0 | 0 | 25 | 1 | 0 | 0 | 0 | 0 | 3 |
| immune | 0 | 0 | 0 | 0 | 33 | 0 | 0 | 1 | 0 | 17 |
| granulosa | 0 | 0 | 0 | 0 | 9 | 0 | 2 | 2 | 42 | 1 |
| endothelial | 0 | 0 | 0 | 0 | 4 | 7 | 0 | 0 | 0 | 131 |
| perivascular | 12 | 0 | 0 | 0 | 14 | 0 | 2 | 21 | 2 | 408 |
| stroma | 5 | 0 | 3 | 0 | 67 | 1 | 33 | 47 | 120 | 5,155 |

Integration with sorted DDX4 Ab+ data

Integrated cluster ID

| | mitotic FGCs | RA resp. FGCs | meiotic FGCs | Oo-genesis | NA | endo-thelial | granulosa (w7-10) | granulosa (w10-20) | granulosa (w20-26) | DDX4 Ab+ (5,479 cells) |
|---|---|---|---|---|---|---|---|---|---|---|
| mitotic FGCs | 409 | 1 | 2 | 0 | 31 | 0 | 0 | 25 | 10 | 1 |
| RA responsive FGC | 2 | 92 | 1 | 1 | 13 | 0 | 0 | 0 | 0 | 0 |
| meiotic FGCs | 0 | 1 | 89 | 7 | 26 | 0 | 0 | 0 | 0 | 0 |
| oocytes/oogonia | 0 | 0 | 0 | 23 | 1 | 0 | 0 | 0 | 0 | 2 |
| immune | 0 | 0 | 0 | 0 | 32 | 0 | 0 | 1 | 0 | 40 |
| granulosa | 1 | 0 | 1 | 0 | 30 | 1 | 14 | 3 | 111 | 124 |
| endothelial | 0 | 0 | 0 | 0 | 4 | 7 | 0 | 0 | 0 | 50 |
| perivascular | 32 | 0 | 0 | 0 | 29 | 0 | 0 | 38 | 24 | 4,495 |
| stroma | 1 | 0 | 0 | 0 | 14 | 0 | 23 | 4 | 19 | 767 |

Integration with sorted DDX4 Ab- data

Integrated cluster ID

| | mitotic FGCs | RA resp. FGCs | meiotic FGCs | Oo-genesis | NA | endo-thelial | granulosa (w7-10) | granulosa (w10-20) | granulosa (w20-26) | DDX4 Ab- (6,690 cells) |
|---|---|---|---|---|---|---|---|---|---|---|
| mitotic FGCs | 437 | 2 | 2 | 0 | 24 | 0 | 1 | 0 | 0 | 2 |
| RA responsive FGC | 0 | 90 | 0 | 0 | 10 | 0 | 0 | 0 | 0 | 0 |
| meiotic FGCs | 1 | 2 | 88 | 8 | 18 | 0 | 0 | 0 | 0 | 0 |
| oocytes/oogonia | 0 | 0 | 0 | 23 | 1 | 0 | 0 | 0 | 0 | 21 |
| immune | 0 | 0 | 0 | 0 | 34 | 0 | 0 | 1 | 0 | 74 |
| granulosa | 0 | 0 | 2 | 0 | 30 | 0 | 3 | 17 | 79 | 244 |
| endothelial | 0 | 0 | 0 | 0 | 4 | 7 | 0 | 0 | 0 | 188 |
| perivascular | 0 | 0 | 0 | 0 | 0 | 0 | 0 | 0 | 0 | 108 |
| stroma | 7 | 0 | 1 | 0 | 59 | 1 | 33 | 53 | 85 | 6,053 |

Our adult ovarian cortex datasets were integrated with a publicly available fetal ovary scRNA-seq dataset [18] in order to find any adult cells that would cluster with pre-meiotic germ cells. Integration was carried out sample by sample in order to avoid bias caused by disproportionate cell numbers between the datasets. Numbers of cells contributing to the integrated clusters from fetal and adult samples are shown for each comparison. Integration with unsorted C-sec sample is displayed in main Figure 5. Adult cells clustering with pre-meiotic germ cells are highlighted in red.
